# Transcriptomic analyses reveal groups of co-expressed, syntenic lncRNAs in four species of the genus *Caenorhabditis*

**DOI:** 10.1101/411561

**Authors:** Cinta Pegueroles, Susana Iraola-Guzmán, Uciel P. Chorostecki, Ewa Ksiezopolska, Ester Saus, Toni Gabaldón

## Abstract

Long non-coding RNAs (lncRNAs) are a heterogeneous class of genes that do not code for proteins. Since lncRNAs (or a fraction thereof) are expected to be functional, many efforts have been dedicated to catalog lncRNAs in numerous organisms, but our knowledge of lncRNAs in non vertebrate species remains very limited. Here, we annotated lncRNAs using transcriptomic data from the same larval stage of four *Caenorhabditis* species. The number of annotated lncRNAs in self-fertile nematodes was lower than in out-crossing species. We used a combination of approaches to identify putatively homologous lncRNAs: synteny, sequence conservation, and structural conservation. We classified a total of 1,532 out of 7,635 genes from the four species into families of lncRNAs with conserved synteny and expression at the larval stage, suggesting that a large fraction of the predicted lncRNAs may be species specific. Despite both sequence and local secondary structure seem to be poorly conserved, sequences within families frequently shared BLASTn hits and short sequence motifs, which were more likely to be unpaired in the predicted structures. We provide the first multi-species catalog of lncRNAs in nematodes and identify groups of lncRNAs with conserved synteny and expression, that share exposed motifs.

## 1. Introduction

Long non-coding RNAs are a heterogeneous class of genes which basically comprises transcripts that are predicted to be non-coding and longer than 200nt. The fact that the first described lncRNAs were functional (e.g. XIST^1^, HG19^2^), coupled to the drastic decrease in price of the high-throughput sequencing technologies, promoted genome-wide screenings for lncRNAs in several vertebrate genomes. Only in the human genome there are currently annotated roughly 60K lncRNAs^3^. However, our knowledge of lncRNAs in non vertebrate species is very limited, with insects being the only exception. Indeed, several recent studies have annotated lncRNAs in several insect genera, including *Drosophila*^*4,5*^, *Anopheles*^*6*^, *Aedes*^*7*^, Apis^8^, *Polistes*^*9*^, *Plutella*^*10*^, *Tribolium*^*8*^, *Nasonia*^*8*^. and *Bombyx*^*11*^. In the phylum Nematoda only one species has been screened so far (*Caenorhabditis elegans*)^12,13^.

Little is known about the functionality of most lncRNA annotated to date, despite great efforts to characterize them. Most studies agree that expression of lncRNAs is often specific to certain tissues, cell types or developmental stages, and that their expression and sequence conservation are lower when compared to those of protein-coding genes^14,15^. In fact, lncRNAs seem to have a fast evolutionary turnover rate and therefore a large fraction of annotated lncRNAs may be species specific^16^. Understanding the evolutionary conservation of lncRNAs is challenging. On the one hand, overall sequence conservation is low despite some patches of sequence are highly conserved, which may be related with their functionality^6,17,18^. On the other hand, secondary structure seems to be important to maintain the function, as some lncRNAs with highly divergent sequences are known to conserve structure and therefore functionality^19,20^. Whether lncRNAs are overall highly structured is still unclear; some studies suggest that they might be less structured than protein-coding genes (at least *in vitro*)^21^ while others suggest that they seem to be highly structured *in vivo*^*22*^. Importantly, a previous study found footprints of selection related with secondary structure, suggesting that the structural conformation may be selectively maintained at least in a fraction of lncRNAs^18^. In addition, experimentally-determined secondary structures are often not available for lncRNAs, and *in silico* predictions may not be highly accurate due to the difficulty of modeling long structures^23^. As a result, detecting homology between lncRNAs in different species is challenging.

To date several studies have aimed to identify lncRNA homology relationships in both vertebrate and insects. Most studies inferred homology relationships between lncRNAs predicted from RNAseq data from different species using sequence similarity (BLASTn or multiple sequence alignments), despite known limitations^24–27^. For instance, a previous study showed that more than 70% of the lncRNAs cannot be traced to homologs in species that diverged more than 50Myr ago^25^. Trying to predict lncRNA homologs extrapolating the coordinates in genome-wide alignments is a weak strategy, first because of the low overall conservation between species, and second because genomic sequence conservation does not necessarily imply conserved transcription^25^. Other authors suggested that homology relationships may be merely based on structural conservation. For instance, Toraninsson *et al* (2006) compared human and mouse genomes and detected regions with shared expression that had a conserved secondary structure despite a lack of sequence conservation^28^. In a more recent study, Seemann *et al*^*19*^ experimentally validated human-mouse lncRNA pairs with shared expression and structure despite their low abundance and low sequence similarity^19^. Another recent study failed to identify structure conservation in three well studied functional lncRNAs: HOTAIR, SRA and Xist^29^, but lately other authors detected structure conservation (in HOTAIR and RepA) by increasing alignment depth, using a sliding window approach and a more sensitive statistical metric^30^. Finally, others tried to identify orthologs using broader strategies, for example using microhomology, experimentally validated secondary structures and, in some cases, synteny^31,32^. However, and due to the complexity of this pipeline, it has so far only been applied to individual lncRNAs (e.g. roX, HOTAIR and COOLAIR).

We here set out to annotate lncRNAs in four different species of nematodes of the genus *Caenorhabditis* and study their evolutionary relationships. *Caenorhabditis remanei* and *C. brenneri* are out-crossing species while *C. elegans and C. briggsae* are hermaphroditic species, which evolved independently from out-crossing ancestors. Of note, out-crossing species are hyperdiverse (e.g. *C. brenneri* carries the highest molecular diversity described in any eukaryote^33^) while hermaphroditic species have very low levels of genetic polymorphism (e.g. *C. elegans* have lower genetic diversity levels than human^34^). In addition, divergence levels between *Caenorhabditis* species are quite high. An early study showed that species within *Caenorhabditis* are as genetically divergent as different orders in tetrapod classes^35^. In addition, synonymous-site divergence for *Caenorhabditis* species is saturated^36,37^. Synonymous substitution rate (ds) for *C. elegans* (0.52^36^) is much higher than for human (0.18^38^). However, synteny seems to be quite conserved among *Caenorhabditis* species, particularly for the X chromosome^39^. Given the high divergence between the studied species, we performed a large scale analysis to identify homology between lncRNAs annotated from RNAseq data using a combination of approaches: syntenic relationships, sequence conservation and structural conservation.

## 2. Materials and methods

### 2.1. Worm Propagation

Our study focused on four different *Caenorhabditis* worm species: *C. elegans* (N2 strain), *C. briggsae* (AF16), *C. remanei* (PB4641) and *C. brenneri* (PB2801). Worms were propagated on solid media in nematode growth medium (NGM) or in liquid S-Basal media with OP50.1 *E. coli* bacteria as food source at 20°C. Worms were synchronized by bleaching, L1 worms were plated at 500 worms per plate, and harvested at L4-Adult stage (3 days growth). In liquid media, L1 worms were grown to late L4/early adult stage in 250 ml Erlenmeyer flasks at worm densities 1000 worms / ml in 50 ml S-Basal media containing OP50.1 bacteria at an initial OD600 = 1.0 with shaking of 100rpm. Worms were harvested off plates or pelleted by centrifugation, washed 4 times with ice-cold M9 buffer by centrifugation at 200g for 1 min. After final centrifugation, M9 buffer was aspirated to the top of the worm pellet, and then a 4X volume of Trizol Reagent (ref. 15596026, ThermoFisher Scientific) was added and worms were flash-frozen in liquid nitrogen and stored at -80°C until RNA extraction.

### 2.2. Total RNA extraction and polyA selection

For total RNA extraction, samples underwent four rounds of freezing in liquid nitrogen, thawing in a 37°C water bath and vortexing for 1 min. After 5 min at room temperature, 180 µl of chloroform were added, samples were shaked vigorously for 15 sec to mix and then were incubated at room temperature for 2 min. After a centrifugation at 12000g, for 15 min at 4°C, the aqueous upper phase was transferred into a new tube, 500 µl of isopropanol were added and samples were frozen at -80°C for 20 min. Then, samples were centrifuged at 12,000g, for 15 min at 4°C to pellet the DNA, the supernatant was removed and discarded, and a final cleaning with 1 ml of cold 70% ethanol was done (centrifugation at 12,000g, for 10 min at 4°C). Finally, after removing the supernatant and leaving the tubes with the lid open to dry the pellet, 30 µl of RNase-free water were added to resuspend the samples. RNA samples were treated with DNase I (Invitrogen™, Carlsbad, California) following the manufacturer’s guidelines to remove traces of Genomic DNA. Total RNA integrity and quantity of the samples were assessed using the Agilent 2100 bioanalyzer with the RNA 6000 Nano LabChip Kit (ref. 5067-1511, Agilent,) and NanoDrop 1000 Spectrophotometer (Thermo Scientific). For three samples (*C. elegans, C. remanei* and *C. brenneri*) we obtained PolyA+ RNA after purifying total RNA by two rounds of selection using MicroPoly(A)Purist™ Kit according to manufacturer’s instructions (ref. AM1919, Ambion), and the quality of the samples was controlled as above.

### 2.3. Library preparation and sequencing

Libraries were prepared using the TruSeq Stranded mRNA Sample Prep Kit v2 (ref. RS-122-2101/2, Illumina) according to the manufacturer’s protocol for all samples. All reagents subsequently mentioned are from the TruSeq Stranded mRNA Sample Prep Kit v2 if not specified otherwise. For three samples, total RNA (Total_ *C. elegans*, Total_*C. briggsae* and Total_*C. remanei*) was used to start the library preparation to check if they were differences between polyA+ RNA selection methods. For these samples, 1 µg of total RNA were used for poly(A)-mRNA selection using streptavidin-coated magnetic beads. This initial step was skipped in the already selected polyA+ RNA samples (see above), for which 100 ng of the samples were used to start the library preparation. To assess the lower limit of detection of gene expression in these experiments, 1 µl of 1:10 dilution of Spike-In Mix 1 from ERCC RNA Spike-In Mix (ref. 4456740, Life Technologies) was added in two of the polyA+ RNA samples (*C.elegans and C.remanei*) before starting the library preparation. Briefly, all samples were subsequently fragmented to approximately 300bp. cDNA was synthesized using reverse transcriptase (SuperScript II, ref. 18064-014, Invitrogen) and random primers. The second strand of the cDNA incorporated dUTP in place of dTTP. Double-stranded DNA was further used for library preparation. dsDNA was subjected to A-tailing and ligation of the barcoded Truseq adapters. All purification steps were performed using AMPure XP Beads (ref. A63881, Agencourt). Library amplification was performed by PCR on the size selected fragments using the primer cocktail supplied in the kit. Final libraries were analyzed using Agilent DNA 1000 chip (5067-1504, Agilent) to estimate the quantity and check size distribution, and were then quantified by qPCR using the KAPA Library Quantification Kit (ref. KK4835, KapaBiosystems) prior to amplification with Illumina’s cBot. Libraries were loaded and sequenced 2 x 50 on Illumina’s HiSeq 2000/2500.

### 2.4. Transcriptome assembly and d*e novo* annotation of intergenic lncRNA in nematodes

We used RNAseq data to annotate lncRNAs *de novo*. In particular, we obtained stranded paired-end, 50bp-long read RNAseq data from *C. elegans* (2 biological replicas, 75 and 107 My reads each), *C. briggsae* (79 My reads), *C. remanei* (2 biological replicas, 103 and 110 My reads each) and *C. brenneri* (83 My reads). In those species having two biological replicas we added spike-ins to one of the libraries to evaluate the accuracy of the RNAseq experiment. The quality of the reads was assessed with FastQC^40^. No extra filtering was needed since the overall quality was high (average Phred quality score >34 in all samples) and no adapters were detected in the reads provided by the sequencing unit. We aligned reads to the reference genomes for each species available in WormBase version WS256 with TopHat v2.0.9^41^, which uses Bowtie v2.1.0.0 in the first step^42^. Then, we assembled aligned reads into individual transcripts with Cufflinks v2.2.1^43^. Finally, we used the utility Cuffcompare included in Cufflinks to annotate our transcript assemblies using a reference annotation, which helps to sort out new genes from those previously annotated. In those species with biological replicas we obtained a merged transcriptome using Cuffmerge, and we finally used Cuffquant and Cuffnorm to quantify the expression (in FPKM) of the reconstructed transcripts normalizing by library size. In addition, we used spike-ins to compare the estimated gene expression versus the expected provided by the company. The accuracy of the experiment was high since the correlation between the expected amount of spike-ins and the estimated in FPKM was >0.98 in all cases (**Supplementary Fig. S1**). The reproducibility of the experiment was also high, since the correlation of the estimated FPKM values for spike-ins between species was 0.99 (**Supplementary Fig. S1**). Thus, no extra filtering was performed to the reconstructed transcripts. To annotate lncRNA we first collected all reconstructed transcripts classified as lincRNA (only in *C. elegans*), ncRNA and unannotated, we removed those overlapping with protein-coding genes annotated in WS256 using BEDOPS tools^44^ and we discarded transcripts shorter than 200 ncl (**Supplementary Table S1**). Finally, we evaluated the coding potential of the remaining transcripts using CPC v0.9^45^ to obtain the final list of lncRNA candidate genes (1,059 in *C. briggsae*, 1,765 in *C. remanei*, 3,285 in *C. brenneri* and 1,526 in *C. elegans*, **Supplementary Table S1**).

### 2.5. Validation of lncRNA expression by semi-quantitative RT-PCR

RNA isolated from N2 strain and free of genomic DNA was reverse-transcribed into cDNA using SuperScript® II Reverse Transcriptase (Invitrogen™), including a minus reverse transcriptase control. Semi-quantitative RT-PCR was performed in a DNA engine tetrad 2 thermal cycler (Bio-Rad) with Taq Mix (Donsheng Biotech) and a final primer concentration of 0.5µM. The primers used in this study are listed in **Supplementary Table S2**. ‘Touch-down’ PCR amplification conditions were as follows: PCR started with denaturing step at 95°C for 2 min, followed by 15 cycles of touchdown amplification. Every cycle consisted in 3 steps, each for 30 seconds: denaturation at 95 °C, annealing (starting with a temperature 7°C higher than annealing temperature and decreasing -0.5°C per cycle until annealing temperature of primers), and an extension at 72°C. Then, 20 additional cycles were carried out at annealing temperature (57.3°C, 60.2 °C, 63.6 °C, 60.2 °C for XLOC_040084, XLOC_005215, XLOC_007633, and XLOC_036972, respectively). Final primer extension was performed at 72°C for 5 minutes. All PCR and RT-PCR products were visualized in 1.5% agarose gels.

### 2.6. Classification of lncRNAs into blast-based families

We first performed BLASTn searches for pairs of species using an e-value cut-off of 1e-3. We then selected the reciprocal hits and we finally classified lncRNAs from the four different species into families using the in-house script classifyFamiliesv5_VennGH.py (see below).

### 2.7. Classification of lncRNA into secondary structure based families

To evaluate the presence of conserved structural elements within lncRNAs we used the software BEAGLE^46^. This software compares secondary structures encoded using the BEAR notation by performing pairwise global or local alignments and provides measures of structural similarity and statistical significance for each alignment. We used this software to compare the secondary structures of lncRNAs, intergenic regions and rRNAs, the two later used as negative and positive controls respectively. We first computed the secondary structures individually for each sequence using the RNAfold software from the Vienna Package 2.0 with default parameters ^47^. The secondary structures computed with RNAfold were used as input for beagle. We ran beagle for all possible pairwise comparisons using local alignments since it has been observed that the function of some lncRNA is restricted to small regions with conserved secondary structure. We finally used the classifyFamiliesv5_VennGH.py script to classify sequences with significant matches (Zscore >3) into families (see below). Intergenic and rRNA sequences were obtained from Wormbase WS256. We ran the pipeline for the 102 rRNA sequences retrieved. For intergenic regions, for each of the 4 species we randomly selected 250 sequences between 200-1500 ncl, we discarded those containing candidate lncRNA and then we ran RNAfold and beagle as previously explained. We ran the pipeline for intergenic regions 50 times.

### 2.8. Classification of protein-coding genes and lncRNA into syntenic families

We further classified the candidate lncRNA into families based on syntenic relationship using an in-house python pipeline available in GitHub (https://github.com/Gabaldonlab/projects/tree/master/syntenic_families). First of all, we created a file including orthology relationships for all genes annotated in the four species using the wormbase_orthoParalogsGH.py script. Subsequently, we used the script synteny_nematodesv4GH.py to obtain a file including pairwise syntenic relationships between lncRNA from the four studied species. Briefly, the script compares the genomic context of lncRNA between two different species. The user can specify the number of genes that will consider at the left and the right sides of each lncRNA, the overall minimum of shared genes and the minimum number of shared genes at each side of the lncRNA. We ran this script considering three genes at each side of a given lncRNA, a minimum of overall 3 shared genes and a minimum of one shared gene in each side of a given lncRNA. Finally, we classified lncRNA from the four different species into families using the script classifyFamiliesv5_VennGH.py.

To validate the performance of our pipeline we analysed 1,000 protein-coding orthologous families randomly selected. We classified 50.3% of the selected genes, and in consequence the number of genes classified per syntenic family was significantly smaller compared to the orthologous families (median= 4 and 5 respectively, p-value =1.7e-80). The number of genes in a given syntenic family correctly assigned is significantly higher than the not correctly assigned (median = 3 and 0 respectively, p-value= 2.4e-217). 19.8% of the syntenic families include genes from more than one orthologous family; in those cases we discarded the orthologous family having the lower number of shared genes. Overall, the number of genes from the matched orthologous family not assigned is quite low (median=1) and significantly smaller than the number of genes per orthologous family (median = 1 and 5 respectively, p-value=1.7e-135). Overall the percentage of genes correctly assigned per syntenic family is very high (median=100%, mean= 92.1%) and most of the genes for each orthologous family were classified in a given syntenic family (median =75%, mean=66.7%). Altogether these results indicates that the rate of misclassification using our pipeline is very low but also that roughly half of the genes could not be classified using our syntenic approach.

### 2.9. Sequence conservation and motifs

We also used a blast strategy to identify syntenic families with some degree of sequence conservation. We performed a BLASTn search of all annotated genes against themselves and we discarded those genes with no hit using a relaxed e-value of 1e-3. Motif identification was performed using the MEME suit (version 4.11.4) ^47^, using windows ranging from 10-40 ncl and an e-value threshold of 0.1. We searched for motifs in each of the syntenic families and for 1,000 randomly generated families as a control. Random families are comprised of sequences belonging to different syntenic families that may belong or not to the same species.

### 2.10. Identification of conserved structures within lncRNAs

Since secondary structure is difficult to predict for long sequences, we split the sequences in overlapping windows of 80ncl + 40 step and 200ncl +100 step, and repeated the analyses described above (RNAfold + Beagle, detailed in section 2.6). We finally calculated the accessibility per position using the RNAplfold from the Vienna Package 2.0, using a window length of 80ncl and -u4 and -u10 options.

## 3. Results and discussion

### 3.1. De novo annotation of intergenic lncRNA in nematodes

Our study aimed to *de novo* annotate lncRNAs in four different *Caenorhabditis* worm species (*C. elegans,* N2 strain; *C. briggsae* AF16; *C. remanei* PB4641; and *C. brenneri* PB2801) and to establish homology relationships between them. Thus, the first step consisted of obtaining high quality RNAseq data. Since lncRNAs are expressed in a stage-dependent manner^12^, worms were synchronized and harvested at the same developmental stage (see material and methods for details). We followed an experimental protocol to obtain stranded paired-end, 50bp-long read RNAseq data from the four studied species. Subsequently, we applied a bioinformatics pipeline to annotate candidate lncRNAs after assessing the accuracy and the reproducibility of the experiment using spike-ins from reference molecules (see material and methods). After discarding candidate transcripts overlapping protein-coding genes, shorter than 200 nucleotides, or showing signals of coding potential, we obtained the final list of candidate lncRNA genes: 1,059 in *C. briggsae*, 1,765 in *C. remanei*, 3,285 in *C. brenneri* and 1,526 in *C. elegans* (**Supplementary Table S1**, see material and methods for details). We next checked whether any lncRNAs in our data set was known to be functional. There are currently four experimentally-validated functional lncRNAs in *C. elegans* according to lncRNAdb^48^. Of these, *Linc-3* and *Rncs-1* genes are included in our catalog but *YRNAs* and *7sk* genes were filtered by our pipeline, as they are both shorter than 200ncl. We finally selected four lncRNAs (XLOC_040084, XLOC_005215, XLOC_007633, and XLOC_036972) to confirm the expression of their transcripts at stage L4/adult in wild type strain N2. In brief, we isolated RNA, reverse-transcribed it into cDNA, and amplified target regions of the transcripts along with a positive control of genomic DNA and a negative control of minus reverse transcriptase. Results were resolved in a 1.5% agarose gel. As shown in **Supplementary Fig. S2**, all four fragments were amplified by RT-PCR confirming the expression observed by RNA-sequencing.

The largest number of candidate lncRNAs were annotated in the two out-crossing species, especially in *C. brenneri* (**Supplementary Table S1**). We tested whether the polyA enrichment performed in different steps of the protocol and using different kits (see material and methods) may influence the annotation of lncRNAs. For this, we compared the expression values of all the annotated lncRNAs in the two species that were sequenced using the two protocols (*C. elegans* and *C. remanei*). Genes expression values assessed from data obtained the two protocols were highly correlated (r^2^=0.92 for *C. elegans*, r^2^=0.90 for *C. remanei,* **Supplementary Fig. S3**) and therefore the two library preparation methods do not seem to bias the gene expression quantification. In agreement with our finding, the number of annotated protein-coding genes in their reference genomes is also higher in *C. remanei* and *C. brenneri* (31,436 and 30,660 respectively in WS256) than in *C. elegans* and *C. briggsae* (20,251 and 22,504 respectively in WS256). In *Caenorhabditis* species the genome size of self-fertile nematodes such as *C. elegans* and *C. briggsae* is 20-40% smaller than out-crossing species such as *C. remanei* and *C. brenneri*^*39*^. This reduction seems to be caused by patterns of gene losses affecting different gene types in roughly similar proportions^39^. This may explain why in the out-crossing species we annotated more lncRNAs than in the hermaphroditic ones.

We compared GC content, length, number of exons and expression levels for the annotated lncRNAs and protein-coding genes in the four studied species. Overall, protein-coding genes are longer, have a larger number of exons, have higher GC content and are expressed at higher levels than lncRNAs and, in all cases, differences are significant (p-value<2.2e-16 for all comparisons, **Fig. 1**). Thus, our results are in agreement with previous findings in other species^8,49^ and suggest that, despite the radically different sexual behaviours in the studied species, the major differences are protein-coding-lncRNA instead of species driven.

**Figure 1:**
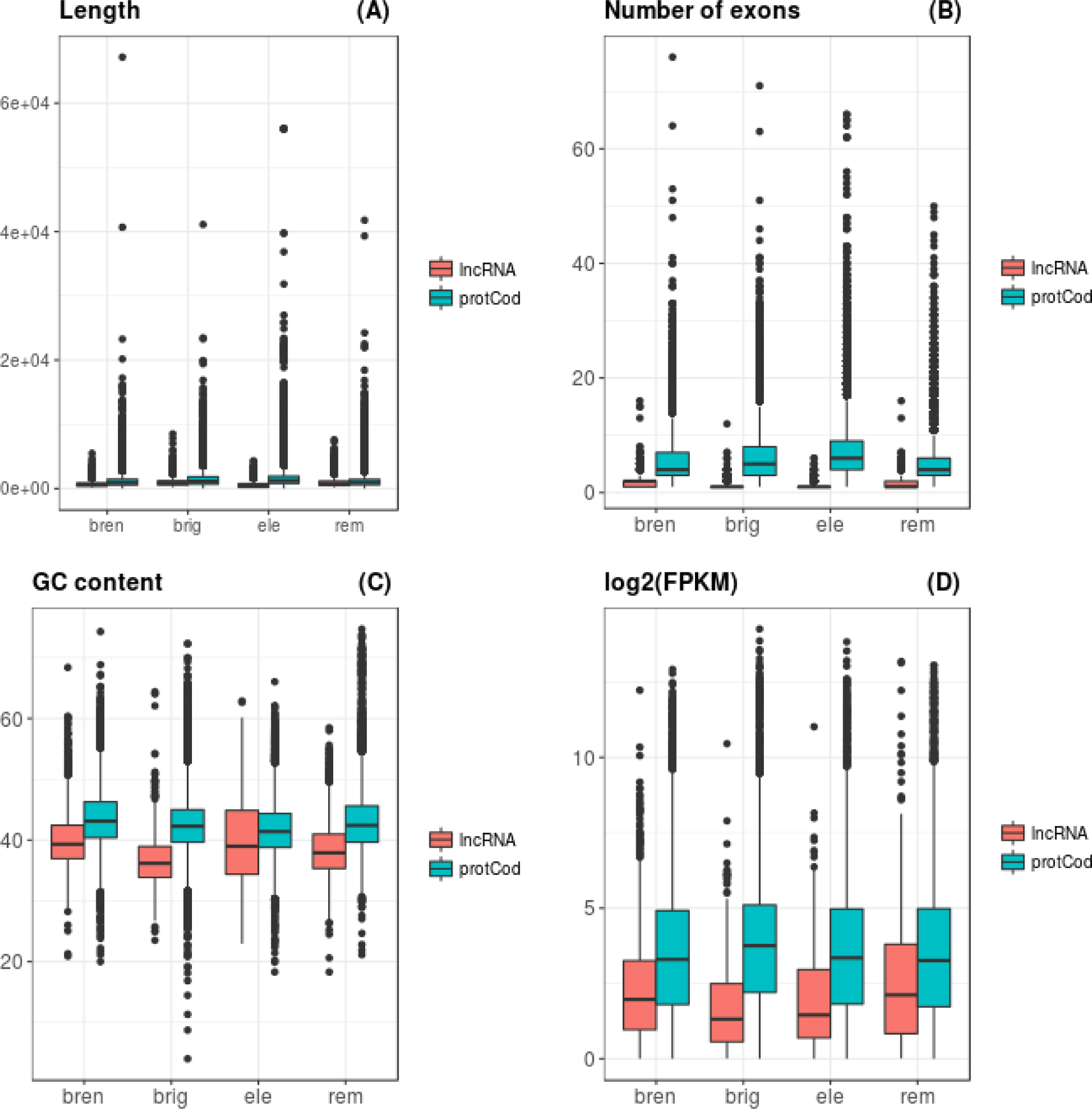
Box-plots showing length **(A)**, number of exons **(B),** GC content **(C),** and expression values **(D)** for the annotated protein-coding and lncRNA genes in all studied species. In Figure 2D we discarded genes with log2(FPKM)<0.

### 3.2. Synteny as a proxy to identify homologous lncRNAs

Establishing homology relationships among lncRNAs is challenging due to their low sequence conservation. Even more challenging is the establishment of homology relationships between lncRNAs of species as divergent as the nematodes considered here. Nevertheless, we made a first attempt to classify genes using a reciprocal BLASTn hits approach. To do so, we first performed BLASTn searches for pairs of species using an e-value cut-off of 1e-3. We then selected the reciprocal hits and we finally classified lncRNAs from the four different species into clusters of co-expressed genes using an in-house script (see material and methods, **Supplementary Table S3**). As expected, the number of genes classified into families was very low (204 out of 7635, **Fig. 2A**). Most of the families (64) comprised transcripts from merely two species, 12 families included transcripts from three species and no families included transcripts from the four species (**Fig. 2A**).

**Figure 2:**
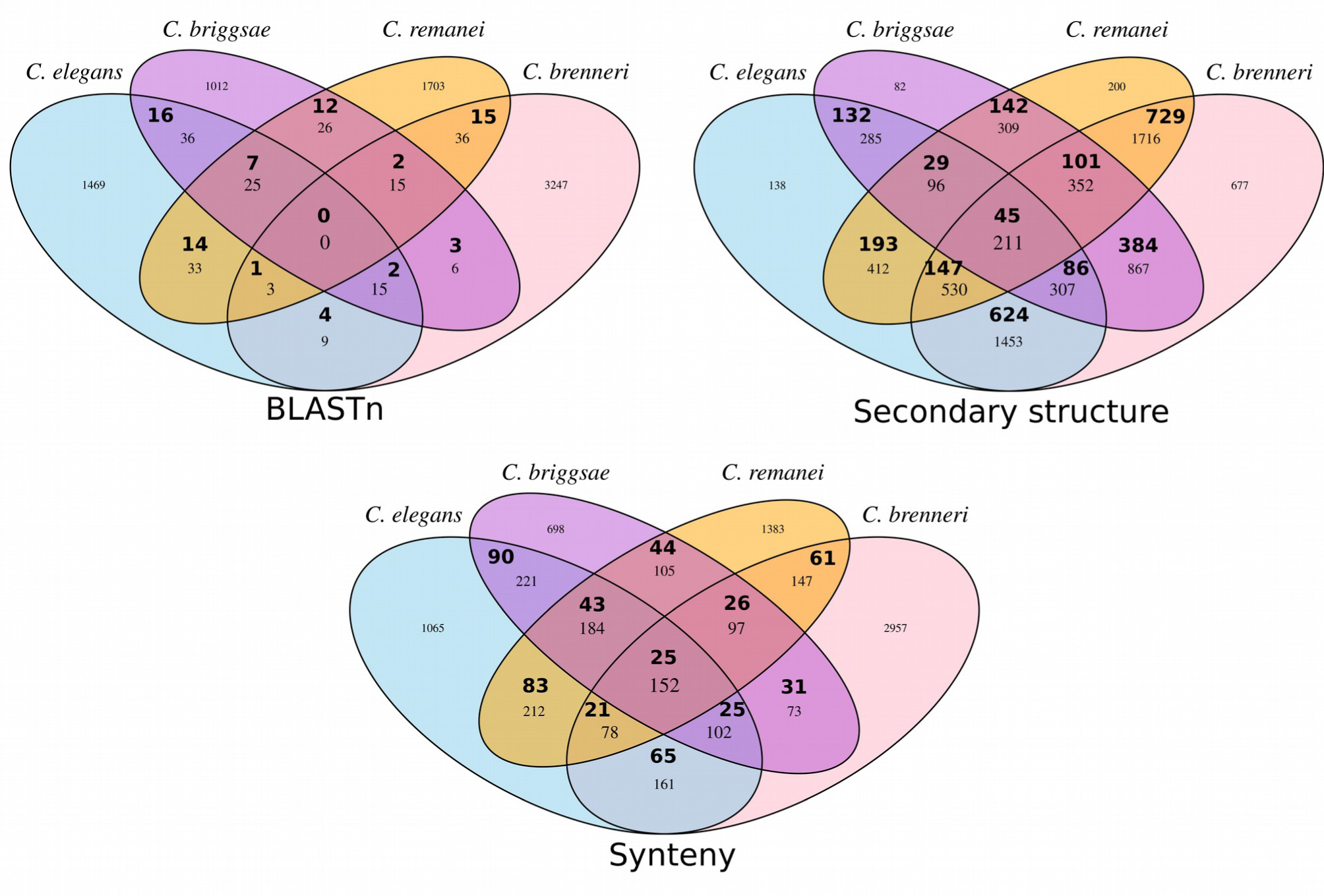
Venn diagram showing the number of lncRNA families (in bold) and genes for the blast (**A**), secondary structure (**B**) and syntenic classification (**C**). *C. elegans* in blue, *C. briggsae* in dark pink, *C. remanei* in orange and *C. brenneri* in light pink.

Knowing that selection may act on conservation of the secondary structure rather than on sequence, we analyzed the patterns of structural conservation of lncRNAs. Since lncRNAs are known to be weakly conserved at the sequence level, we used the software BEAGLE^46^ to study the secondary structure similarity of lncRNAs, since this algorithm compares structures without requiring a sequence alignment. The software provides an associated z-score for each comparison, and a z-score higher than three is indicative of significant similarities. We classified transcripts with significant matches (Zscore =>3) into families (**Supplementary Table S4**). Most of the genes were classified into families (6,538 out of 7,635 genes, **Fig. 2B**). The majority of the families (2,204) comprised genes from two species, 363 families included transcripts from three species and 45 families included transcripts from the four species (**Fig. 2B**). Secondary structure is difficult to predict for long sequences as lncRNAs^23^, and therefore predictions should be considered with caution.

We also used a syntenic approach to classify lncRNAs into clusters of co-expressed syntenic genes, referred hereafter as syntenic families. Syntenic families include genes from the different species that share the same genomic context, meaning that they are surrounded by orthologous genes. Therefore, genes within each family are likely to share orthology relationships. To do so, we implemented a custom pipeline (see Materials and Methods). We benchmarked this pipeline with randomly-chosen 1000 protein-coding orthologous families, containing 1,000 *C. elegans*, 1,232 *C. briggsae*, 1,870 *C. remanei* and 1,646 *C. brenneri* genes. Our pipeline classified a total of 2,895 orthologous genes in syntenic families (61.2% in *C. elegans, 56.9*% in *C. brigssae, 47.4*% in *C. brenneri and 42.9*% in *C. remanei*), meaning that our pipeline classified roughly half of the genes. We assessed the correspondence between syntenic relationships and the known orthology relationships of the protein-coding genes. The positive predictive value is 73%, meaning that synteny-based orthology assignments are very likely to be correct. Importantly, 80.2% of the syntenic families include genes from a single orthologous family, meaning that syntenic families are not likely to be fragmented. We evaluated whether the chance of detecting syntenic families is affected by the phylogenetic distance, we compared the number of classified syntenic families including any of the two most closely related species (*C. briggsae* and *C. remanei*) (**Supplementary Table S5**). The number of families does not drop when increasing the phylogenetic distance, suggesting that *Caenorhabiditis* genomes are highly reshuffled even for closely related species. However, other factors may affect the chance of detecting syntenic families. For instance, *C. briggsae* and *C. elegans* are the only two species in which their genomes are assembled in six chromosomes and this completeness of the genome may favor the detection of syntenic families. Altogether, we can use our conservative method as a proxy to detect homology relationships between lncRNAs, even in the absence of sequence conservation (see Materials and Methods for more details)

The number of syntenic families and lncRNAs classified per species is shown in **Fig. 2C** and **Supplementary Table S6**. We classified 20% of the lncRNAs (1,532 out of 7,635) into syntenic families, which is a much lower rate than for protein-coding genes (50.3%). This result supports the idea that most lncRNAs may be species specific and that lncRNA families have a rapid turnover^6,16^. Similar to protein-coding genes, the number of syntenic families is independent of the phylogenetic distance (**Supplementary Table S5**). Of note, 25 families contains genes from the four studied species, suggesting that their functions during development may be conserved in *Caenorhabditis*. Interestingly, the functional lncRNA *Rncs-1* is included in the syntenic family 146, which also includes two *C. remanei* genes.

### 3.3. Identification of conserved motifs across syntenic families

As most lncRNAs exhibit weak or untraceable primary sequence conservation, we used a motif-based search method to identify conserved domains in different species. Using the MEME suite ^47^ we searched for motifs within the syntenic families. We found that 14.6% of the syntenic families (75 out of 514) shared motifs between at least two of the four species (**Supplementary Table S7**). To evaluate whether this number is higher than expected by chance, we randomly generated 1,000 artificial families shuffling genes from different families and searching for motifs. We found that only 3 families (0.3%) shared conserved motifs in these randomized controls, supporting that syntenic families are enriched in conserved sequence motifs. We also used a blast strategy to evaluate the presence of conserved regions. We performed BLASTn searches within the syntenic families using an e-value threshold of 1e-3. Then we computed a blast score for each family, dividing the number of blast hits by the number of pairwise comparisons (including genes from the same species). 13.23% of the syntenic families (68 out of 514) had a minimum score of 0.01 or higher (**Supplementary Table S8**). This number is higher than expected by random, since the score was 0 in all cases when we applied this procedure to randomized families. Importantly, roughly half (35 out of 68) of the families with blast hits also presented conserved motifs. Those are strong candidate genes to be functional, since minor sequence conservation may be sufficient to ensure conserved lncRNAs functionality. For instance, it has been shown that lncRNAs in one species can functionally replace its ortholog in another species despite a very low sequence conservation^31^.

### 3.4. Structural conservation of syntenic lncRNAs

We first compared the secondary structure of lncRNAs, intergenic and ribosomal sequences, using the two later as negative and positive controls, respectively. Overall structure conservation of lncRNAs is lower than for ribosomal RNAs and similar to intergenic sequences (**Supplementary Fig. S4**). Thus, structural similarities found among lncRNAs are weaker than those expected for rRNAs, which are known to be highly structured. Since previous studies suggest that secondary structure is difficult to predict for long sequences^23^, we also evaluated whether syntenic families are enriched in conserved structures after splitting the sequences in overlapping windows of 80ncl +40 step and 200ncl +100 step. As a control, we randomized sequences from different families. The mean z-score for the syntenic families was lower than three and not significantly different from the randomized sequences, supporting that the secondary structure conservation within families is overall weak. We next assessed, from predicted structures, whether the above described sequence motifs were associated to particular secondary structures. Importantly, we detected that accessibility (meaning the probability of being within unpaired regions) is significantly higher for regions covered with motifs (p-value=4.3e-08 for -u4; p-value=2.3e-08 for -u10; **Supplementary Fig. S5**). Considering this, we hypothesize that these motifs may be involved in binding with other molecules and therefore they may play a role in the functionality of these lncRNAs.

## 4. Concluding remarks

To identify homology within lncRNAs from different species we can use sequence conservation, structural conservation and syntenic relationships. Nematodes are highly divergent at the sequence level and lncRNAs are known to evolve fast, hampering the use of sequence conservation approaches. Instead, we used a synteny-driven strategy that was shown to be an accurate proxy for homology, based on an assessment on protein-coding genes with known orthology relationships. We next refined our results using sequence conservation and structural information. The fraction of lncRNAs classified in syntenic families is lower than for protein-coding genes, suggesting that a large fraction of the predicted lncRNA may be species specific. Genes within families are enriched in conserved motifs which have significant higher accessibility and therefore are more likely to be unpaired. Further studies are needed to elucidate whether these highly accessible motifs conserved in different species are important for their functionality by bounding to other molecules. Finally, we show that the reduction in the genome size of the self-fertile nematodes also affects the lncRNA class, since in these species the number of annotated lncRNA was lower than in out-crossing species. We provide the first catalog of lncRNAs in nematodes including species specific genes, which may be a potential source for novel functions, and genes sharing homology between species, which may be important for their development.

## Supplementary data

Supplementary data are available.

## Acknowledgments and funding

We are very grateful to Eugenio Mattei for kindly providing us a local version of the Beagle software. We also thank Aaron M. New for his help with worm cultures and RNA extraction and Marcos Francisco Pérez for his advice in the validation of lncRNA expression. TG group acknowledges support from the Spanish Ministry of Economy, Industry, and Competitiveness (MEIC) for the EMBL partnership, and grants ‘Centro de Excelencia Severo Ochoa 2013-2017’ SEV-2012-0208, and BFU2015-67107 cofounded by European Regional Development Fund (ERDF); from the CERCA Programme / Generalitat de Catalunya; from the Catalan Research Agency (AGAUR) SGR857, and grant from the European Union’s Horizon 2020 research and innovation programme under the grant agreement ERC-2016-724173 the Marie Sklodowska-Curie grant agreement No H2020-MSCA-ITN-2014-642095.

---

**Supplementary Figure S1:** Expected and estimated amount of spike-ins in log2(FPKM) for *C. elegans* **(A),** *C. remanei* **(B)** and between the two species **(C)**.

**Supplementary Figure S2.** LncRNAs endogenous expression of wild type strain N2 at L4/adult stage determined by semi-quantitative RT-PCR. Agarose gel (1.5%) showing the amplification product of reverse transcribed (RT) lncRNAs: **XLOC_040084** (lane 2-5), **XLOC_005215** (lane 6-9), **XLOC_007633** (lane 10-13), **XLOC_036972** (lane 14-17). **Lane 1:** 1Kb Plus Ladder, **lane 2, 6, 10, 14:** N2 genomic DNA (positive control), **lane 3, 7, 11, 15:** N2 cDNA, **lane 4, 8, 12, 16:** minus reverse transcriptase control (RT negative control), **lane 5, 9, 13, 17:** no-template control (NTC, RT-PCR negative control), **lane 18:** Low Molecular Weight ladder (LMW).

**Supplementary Figure S3:** Expression values of lncRNA (expressed in log2(FPKM)) using the two different protocols for *C. elegans* and *C. remanei*.

**Supplementary Figure S4:** Density plot showing z-scores estimated using Beagle software for random intergenic regions, lncRNA classified in syntenic families and rRNA. Dashed vertical line is located in z-score=3.

**Supplementary Figure S5:** Accessibility values for positions covered or not by motifs using -u10 or -u4 parameters.

**Supplementary Table S1**: Annotation of lncRNA in the four *Caenorhabditis* species.

**Supplementary Table S2**. List of primers used in the study to validate expression of lncRNAs by semi-quantitative RT-PCR. Abbreviations: melting temperature (Tm), nucleotides (nt), base pairs (bp).

**Supplementary Table S3**: Clusters of co-expressed genes with sequence similarity according to the reciprocal BLASTn hits approach used (see material and methods for details).

**Supplementary Table S4:** Clusters of co-expressed genes with secondary structure similarity estimated using BEAGLE software (see material and methods for details).

**Supplementary Table S5:** Number of clusters of co-expressed syntenic genes (proteing coding genes and lncRNA) including any of the two most closely related species (*C. briggsae* and *C. remanei*).

**Supplementary Table S6:** Clusters of co-expressed syntenic genes (see material and methods for details).

**Supplementary Table S7**: Table showing the syntenic families with significant MEME motifs.

**Supplementary Table S8**: Blast scores computed for the syntenic families (see material and methods for details).

